# Domestication of *Campylobacter jejuni* NCTC 11168

**DOI:** 10.1101/591701

**Authors:** Ben Pascoe, Lisa K. Williams, Jessica K. Calland, Guillaume Méric, Matthew D. Hitchings, Myles Dyer, Joseph Ryder, Sophie Shaw, Bruno S. Lopes, Cosmin Chintoan-Uta, Elaine Allan, Ana Vidal, Catherine Fearnley, Paul Everest, Justin A. Pachebat, Tristan A. Cogan, Mark P. Stevens, Thomas J. Humphrey, Thomas S. Wilkinson, Alison J. Cody, Frances M. Colles, Keith A. Jolley, Martin C. J. Maiden, Norval Strachan, Bruce M. Pearson, Dennis Linton, Brendan W. Wren, Julian Parkhill, David J. Kelly, Arnoud H. M. van Vliet, Ken J. Forbes, Samuel K. Sheppard

## Abstract

Reference and type strains of well-known bacteria have been a cornerstone of microbiology research for decades. The sharing of well-characterised isolates among laboratories has parallelised research efforts and enhanced the reproducibility of experiments, leading to a wealth of knowledge about trait variation in different species and the underlying genetics. *Campylobacter jejuni* strain NCTC 11168, deposited at the National Collection of Type Cultures in 1977, has been adopted widely as a reference strain by researchers worldwide and was the first *Campylobacter* for which the complete genome was published (in 2000). In this study, we collected 23 *C. jejuni* NCTC 11168 reference isolates from laboratories across the UK and compared variation in simple laboratory phenotypes with genetic variation in sequenced genomes. Putatively identical isolates identified previously to have aberrant phenotypes varied by up to 281 SNPs (in 15 genes) compared to the most recent reference strain. Isolates also display considerable phenotype variation in motility, morphology, growth at 37°C, invasion of chicken and human cell lines and susceptibility to ampicillin. This study provides evidence of ongoing evolutionary change among *C. jejuni* isolates as they are cultured in different laboratories and highlights the need for careful consideration of genetic variation within laboratory reference strains.

**Impact statement:** In this paper, we comment on the changing role of laboratory reference strains. While the model organism allows basic comparison within and among laboratories, it is important to remember the effect even small differences in isolate genomes can have on the validity and reproducibility of experimental work. We quantify differences in 23 reference *Campylobacter* genomes and compare them with observable differences in common laboratory phenotypes.

**Data summary:** Short read data are archived on the NCBI SRA associated with BioProject accession PRJNA517467 (https://www.ncbi.nlm.nih.gov/bioproject/PRJNA517467).

All assembled genomes are also available on FigShare (doi: 10.6084/m9.figshare.7849268). Phylogeny visualised on microreact: https://microreact.org/project/NCTC11168.

**Repositories:** Short read data are archived on the NCBI SRA repository, associated with BioProject accession PRJNA517467 (https://www.ncbi.nlm.nih.gov/bioproject/PRJNA517467; **Table S1**).

The authors confirm all supporting data, code and protocols have been provided within the article or through supplementary data files.

## Introduction

The sharing of bacterial reference or type strains among laboratories is a fundamental part of microbiology. This often informal, usually uncelebrated, enterprise has supported academic, health, food and veterinary research worldwide underpinning microbiology innovation. The history of the exchange and classification of bacterial type strains has incorporated the work of some of the most influential microbiologists [1]. One such strain belongs to the important food-borne pathogen species *Campylobacter jejuni*.

For *C. jejuni,* the publication of a simplified culturing technique and deposition of a reference isolate at the National Collection of Type Cultures (NCTC 11168) in 1977 (by Martin Skirrow), marked the end of the first century of research into this organism [2]. The first description of an organism likely to be *Campylobacter* was made in Naples in 1884. Theodor Escherich observed spiral bacteria in stool specimens from patients with diarrhoeal disease but he was unable to culture them [3, 4]. Successful isolation of *Bacterium coli commune* (now *Escherichia coli*) from his young dysenteric patients helped pioneer bacterial genetics and lay the foundations of modern microbiology [1, 5]. However, throughout his career, Escherich continued to identify ‘*spirilla’* in cases of cholera-like and dysenteric disease. It is likely that the microorganisms he described were *Campylobacter* with their typical spiral morphology and association with enteritis [4, 6].

Early in the 20^th^ century researchers investigating veterinary cases of foetal abortion and winter dysentery in cattle [7] described several species that would later become part of the *Campylobacter* genus, including *Vibrio jejuni* [8]*, V. fetus* [9]*, V. fetus venerealis and V. fetus intestinalis* [10]. Isolation techniques that permitted the growth of *Campylobacter* from human faeces drew attention to its importance as a human pathogen [11–13]. The genus name *Campylobacter* (meaning curved rod) was proposed by Sebald and Véron in 1963 and subsequently verified in 1973 with the broader acceptance of *Campylobacter* spp. as human pathogens [14, 15]. Skirrow’s more convenient culturing technique and the availability of a model reference strain sparked renewed interest in *Campylobacter* research later in the 20^th^ century [16, 17]. Model strains allowed for comparison of experiments within laboratories and isolates were passed among laboratories across the world [18–23]. When the *C. jejuni* NCTC 11168 genome was sequenced in 2000 [24] this type strain was cemented as an important reference strain for *Campylobacter* research. Additional detail was added to the *C. jejuni* genome following its re-annotation (accession: AL11168.1), including revised coding sequence (CDS) identification incorporating potential for phase variation [25–29].

Today, many aspects of the biology of this organism are well characterised. Identification of genomic regions primed for posttranslational modification, in particular decoration of surface proteins with glycans [30], pseudaminic acid [31–33] and legionaminic acid [34] have improved understanding of the mechanisms of ganglioside mimicry [35], epithelial cell invasion, host immune-evasion, colonisation [36, 37] and development of neurological sequelae such as Guillain-Barré syndrome [38]. Furthermore, insights into virulence traits including strategies to sequester the iron required for infection were detailed using NCTC 11168 [39–41]. Vaccine targets have been identified [42–44] and the mechanisms of core metabolic processes [45, 46], biofilm production [47–51], capsule production [52] and resistance to oxidative stress have been elucidated [53, 54]. Accidental passage through a laboratory worker also identified putative human host adaptations *in vivo* [55].

Since 1977 the NCTC 11168 strain has been an important part of efforts to better understand this pervasive pathogen. However, there are limitations to the use of type strains, the most obvious being that bacteria display considerable variation within species. For example, in *C. jejuni,* some strains cause a significant amount of disease in humans while others do not – owing, in part, to their inability to survive the passage from reservoir host through the food production chain to contaminate human food [56]. This kind of phenotypic variation among strains is well-documented in many species and is a central reason for the growing emphasis on population genomics when trying to understand the ecology and evolution of bacteria [57]. A second, more inconspicuous limitation on the use of type strains shared among laboratories is that they might not all be the same. Strains are not *sensu stricto* clones and may display low levels of genetic variation. Clearly, when frozen there is little opportunity for genome evolution to occur [58]. However, whenever there is growth, for example in the process of sub-culturing isolates, there is an opportunity for genetic variability to be generated within the population. This may be important for interpreting research findings in different groups as even single SNPs can potentially have an impact on phenotype, for example in antimicrobial resistance [59] or host tropism [60]. The aim of our study was to investigate if, over time, multiple passages under potentially different growth conditions in different laboratories have introduced genotypic and phenotypic variation into a collection of NCTC 11168 *C. jejuni*.

## Methods

### Isolates and genome sequencing

Twenty-three laboratory reference *C. jejuni* NCTC 11168 isolates from around the United Kingdom were collected and (re)sequenced. The year in which the laboratory received the isolate is noted along with its known heritage (**Table 1**). DNA was extracted using the QIAamp DNA Mini Kit (QIAGEN, Crawley, UK), according to manufacturer’s instructions and quantified using a Nanodrop spectrophotometer. Genome sequencing was performed on an Illumina MiSeq sequencer using the Nextera XT Library Preparation Kit. Libraries were sequenced using 2 × 300 bp paired end v3 reagent kit (Illumina). Short read paired-end data was trimmed using TRIMMOMATIC (version 0.35; paired-end mode) and assembled using the *de novo* assembly software, SPAdes (version 3.8.0; using the *careful* command). The average number of contigs in the resulting assemblies was 19.7 (range: 13-36) for an average total assembled sequence size of 1,629,408 bp (range: 1,612,402 - 1,694,909 bp). The average N50 contig length was 173,674 bp (range: 100,444 - 271,714 bp) (**Table S1**).

**Table 1:** Summary of genome differences in 23 NCTC11168 isolates.

### Population structure and phylogenies

Sequence alignments and genome content comparison analyses using BLAST were performed gene-by-gene, as implemented in the BIGSdb platform [61, 62] as described in previous *Campylobacter* studies [63–66]. A gene was considered present in a given genome when its sequence aligned to a NCTC 11168 locus with more than 70% sequence identity over at least 50% of sequence length using BLAST [67]. Genomes were aligned by concatenating single-gene alignments using MAFFT [68]. For context, collected NCTC 11168 isolates were augmented with 83 previously published genomes representing the known genetic diversity in *C. jejuni* (**Table S2**). Genes present in 90% or more of the isolate genomes were aligned (1,359,883 bp; **Supplementary File 1**) and a maximum-likelihood phylogeny constructed in FastTree (version 2.1.10; with the generalized time reversible substitution model)[69]. A second alignment of just the collected NCTC 11168 strains was made (1,555,326 bp; **Supplementary File 2**) to build an additional maximum-likelihood tree, which was used as input for ClonalFrame-ML to mask putative recombination sites (version 1.11-3)[70] and visualised in microreact: https://microreact.org/project/NCTC11168 [71].

### Estimating genome variation

Sequence reads were compared to the completed NCTC 11168 reference genome (AL11168.1) using SNIPPY (version 3.2dev)[72] to estimate nucleotide differences between our laboratory reference isolates and the originally sequenced genome. Assembled genomes were annotated with PROKKA (version 1.13)[73] and recombination was inferred using Gubbins (version 2.3.1)[71]. All high performance computation was performed on MRC CLIMB in a CONDA environment [74, 75].

### Phenotype testing

Isolates were recovered from frozen storage on Columbia blood agar (E&O Labs, BonnyBridge, UK) and incubated in microaerobic conditions at 37°C and sub-cultured in Mueller Hinton broth (Oxoid Ltd, Basingstoke, UK) and grown microaerobically overnight at 37°C.

### Bacterial growth assays

Broth cultures were standardised to an OD_600_ nm of 0.05. For growth curves at 37 °C and 42°C, 20 μl of the standardised broth culture was added to 180 μl of Mueller Hinton broth in a microtitre plate. Optical densities were measured at hourly intervals over a period of 48 hours using an OMEGA FLUOstar (BMG LabTech, Aylesbury, UK) plate reader with an atmospheric environment of 10% CO_2_ and 3% O_2_. Growth curve assays were performed in triplicate, with three technical replicates for each biological replicate. Multiple comparisons among isolates at 37°C and 42°C were compared using a one-way ANOVA with a Tukey post-test [76].

### Swarming assays and motility

For each isolate, a 1 ml aliquot of the standardised pre-culture (OD_600_=0.05) was transferred to 5 ml of fresh Mueller Hinton broth and 2 µl pipetted onto the centre of semi-solid Mueller Hinton agar (11.5 g of Muller Hinton Broth, 2.5 g of Agar 3 (Oxoid) in 500 ml of deionised water) and incubated at 42°C for 24 hours. Variation in isolate swarming was observed on Mueller-Hinton motility plates. Motile isolates spread across the plates and halo diameters were measured after 1 day of incubation. Isolates were grouped into three categories: non-motile isolates did not spread across the plate; isolates with halo diameters up to 1.5 cm were categorised as motile; and those with halos of a diameter above 1.5 cm were designated as hyper-motile [36].

### Invasion assays

A chicken gut epithelial cell line (MM-CHiC clone, 8E11; Micromol, Germany) and a human colon epithelial adenocarcinoma cell line (HT29) were used to assay invasion of *Campylobacter in vivo*. A 24-well plate was seeded with 8E11 cells in assay medium (modified McCoy’s 5A/DMEM/F-12 with L-glutamine (5 mM) and supplemented with 5% FBS) and incubated at 37°C in 5% CO_2_ between 4 and 7 days. Liquid cultures were standardised by diluting with Mueller Hinton broth to between 0.030 and 0.080. Aliquots of 200 µl from each isolate were deposited into a 96 well plate and diluted serially. The original stock and dilutions were spread onto Columbia horse blood agar and incubated for 24 hours microaerobically at 42°C. Once the cells had reached confluent growth, the medium was removed and the monolayer washed 3 times with warm PBS. An aliquot of 1 ml pre-warmed antibiotic-free supplemented DMEM medium was added to each well and inoculated with 100 µl 1×10^7^ colony-forming units (CFU). Following incubation in 5% CO_2_ at 37°C for 4 hours, the cells were washed twice with 2 ml PBS supplemented with 4 µl (100 µl/ml) gentamicin and incubated for a further 1.5 hours. Cells were washed 3 times with PBS and an aliquot of 1 ml of warmed TrypLE (Gibco) added to each well and incubated at 37°C for 10 minutes. The lysed monolayer solution was diluted serially and spread onto Columbia horse blood agar in duplicate. Plates were incubated overnight at 42°C in a microaerobic environment and enumerated pre- and post- invasion to calculate the percentage of invaded inoculum. Assays with human HT29 cells were performed with McCoys growth media. Invasion assays were performed in triplicate and analysed using unpaired T-tests with Welch’s correction.

## Results and discussion

### Not all reference strains are equal

Since its deposition at the NCTC there have been two main dissemination hubs of NCTC 11168. Ten of the 23 isolates we collected were obtained by contributing laboratories directly from the NCTC collection, while 13 isolates had come via another laboratory (**Figure 1**). DNA was extracted from each isolate, sequenced, and the genome was assembled (**Table S1**). All 23 isolates clustered closely in the host-generalist ST-21 lineage when compared on a maximum-likelihood phylogenetic tree (**Figure 2A**; https://microreact.org/project/NCTC11168). This suggests that despite some phenotypic heterogeneity, all isolates derived were from a recent common ancestor and no strains were misidentified during passage. Micro-evolutionary differences among closely related NCTC 11168 isolates were observed on a recombination-free phylogeny constructed using ClonalFrameML (**Figure 2B**). Genomes were compared to the original NCTC 11168 genome and as many as 281 SNP differences were observed (up to 15 genes) among collected laboratory strains and the reference (**Figure 2C; Table 1**). Although, in 21 of 23 isolates (91%) there were 32 or fewer SNP differences compared to the reference (**Table 1**). There was an average of 29 SNP differences between the laboratory strains and the reference, and fewest SNPs in any comparison was eight SNP differences (in five genes).

**Figure 1.** The location of laboratories contributing *C. jejuni* NCTC11168 isolates. The most recent NCTC 11168 isolate was obtained by Swansea (isolate 13) in 2016 from the NCTC collection. Other isolates obtained directly from the NCTC collection are coloured black, isolates obtained via a second laboratory are coloured white.

**Figure 2.** Genetic variation among *C. jejuni* NCTC11168 genomes. (**A**) NCTC11168 isolates were contextualised with 83 previously published genomes representing the known genetic diversity in *C. jejuni* (total of 106 isolates). Genes present in 90% or more of the isolate genomes were aligned (1,359,883 bp) and a maximum-likelihood phylogeny constructed in FastTree2 with the generalized time reversible substitution model. The scale bar represents a genetic distance of 0.01. (**B**) Recombination was masked using ClonalFrame-ML to produce an alignment of the NCTC11168 isolates only (n=23; 1,555,326 bp). The scale bar represents 15 nucleotide substitutions. (**C**) The position of all nucleotide substitutions identified using SNIPPY were mapped against the original NCTC11168 genome (AL11168.1). SNPs found within coding regions (CDS) are represented with circles and SNPs located in intergenic regions are represented with an X. Gene names are given where variation was observed in 10 or more of the isolates.

Under ideal storage conditions we might not expect to see any evidence of recent recombination in the laboratory reference strains. Nevertheless, we estimated the number of mutations and recombination events using Gubbins. In total, 436 of the 632 SNPs (69%) we identified were found within protein coding regions, of which 83 were synonymous mutations (19%; **Table 1**). The only isolate where we inferred any recombination was isolate **17**. This isolate has acquired four recombination blocks (combined 14,816 bp, r/m of 9.76) and lost a block of 15 genes (*Cj1319-1333*; **wgMLST supplementary file**), which includes a *maf*-family gene (*maf3/Cj1334*) involved in posttranslational modification of flagellins. Also missing were the *neuC2/Cj1328*, *neuB2/Cj1327*, *ptmA/Cj1332*, and *ptmB/Cj1331* genes involved in the addition of pseuaminic/legionaminic acid to *C. jejuni* flagellins [32, 77, 78]. A knockout mutant of the final gene in this block, *Cj1333*, demonstrated compromised agglutination and reduced invasion (in INT-407 cells)[78]. This region of the *C. jejuni* genome is prone to recombination and has shown a high level of diversity and is often implicated in bacterial virulence [34, 35, 37, 79–82]. Isolate 17 was hyper-motile and also among the most invasive isolates when tested against chicken cell lines, but invaded human cell lines poorly (**Table 2**).

**Table 2:** Summary of phenotype differences in 23 NCTC11168 isolates.

Isolate motility was tested *in vitro* [83] and phenotypic variation was observed among NCTC 11168 isolates (**Table 2**). Since its original dissemination, motile, non-motile and hyper-motile variants have been reported [25, 28, 84]. All three hyper-motile strains were passed between at least two laboratories before entering our collection. Only 50% of the isolates received by laboratories directly from the NCTC collection were motile (**Table 2**). Changes in motility can be a result of differences in the *flaA* and *flaB* genes resulting in attenuated flagella assembly [36]. However, we did not identify any non-synonymous mutations within the *flaA* or *flaB* genes. A shared frameshift mutation was identified in two hyper-motile isolates (11 and 16) within the core motor protein, *fliR* [85–87]. Isolate motility is also influenced by phase-variable gene expression as a result of upstream homopolymeric repeat regions [24, 88, 89]. Several motility associated genes (*maf1/Cj1348*, *maf4/Cj1335* and *maf7/Cj1342c*) were among 31 phase-variable regions recently identified in NCTC 11168 [90] and were among SNPs we identified in non-coding intergenic regions (196 of 632; 31%; **Table 1**). Twelve genes contained nucleotide substitutions in 10 or more NCTC 11168 isolates, of which five have been shown to be subject to phase variation [89]. Growth of motile bacteria in culture media can result in loss of motility as flagella construction is energetically expensive [91, 92]. In batch culture, rapid growth is prioritised and loss of flagella can be advantageous [93, 94].

Adequate flagella construction is an important virulence factor as, in addition to motility, flagella also contribute to invasion and secretion [95, 96], without which colonisation is impaired [28]. The ability of isolates to invade human and chicken intestinal epithelial cell lines was tested *in vitro* by gentamicin protection assay (**Figure 3AB**). Fourteen of twenty one isolates tested invaded the 8E11 chicken cell line more effectively compared to the human HT-29 cell line (**Figure 3C**). Broadly, motile and hyper-motile isolates invaded both cell lines in greater numbers **Figure 3AB**). Several genes containing SNPs in multiple isolates have been shown previously to contribute to increased invasion and virulence, including *mreB*, *cheA, Cj0431, Cj0455, Cj0807* and *Cj1145* [55, 81, 97]. Isolate growth was tested at 37°C and 42°C, with all growing to a higher optical density at avian body temperature (42°C) (**Figure 3D**). Isolate 15 grew particularly poorly at 37°C. We identified the OXA-61 gene in the majority of isolates, but only two were resistant to ampicillin, according to CLSI guidelines (**Isolates 3 and 8; Table 2; Figure 3E**) [98].

**Figure 3.** Phenotype variation among *C. jejuni* NCTC11168 genomes. Invasion assays were carried out for strains categorised by motility phenotypes in (**A**) human HT-29 and (**B**) chicken cell lines. Comparisons were made between (**C**) invasiveness in these cell lines and (**D**) maximum growth at different temperatures. Minimum inhibitory concentration of ampicillin was determined for isolates grouped by source (**E**) and motility (**F**).

### The role of model strains in an age of population genomics

In most cases (21 of 23 isolates; 91%) we observed fewer than 32 SNPs among the laboratory isolate and the type strain deposited in the NCTC archive. However, even these minor changes are associated with observable phenotype differences (motility and invasion as seen here). This could be seen as a challenge to the reproducibility of experiments in different laboratories that use ostensibly identical strains [55, 97]. It is accepted among microbiologists that there is potential for variation among type strains that may display considerable genome plasticity, such as in *Helicobacter pylori* [99]. Consistent with this, variants of *C. jejuni* NCTC 11168 are defined as motile/non-motile, coloniser/non-coloniser for use in specific experiments.

Technical advances in high-throughput genome sequencing and analysis methods continue to improve understanding of *C. jejuni* from bottom-up studies that test the function of specific genes or operons, often with insertion or deletion mutants [55, 97], to top-down comparative genomic approaches in which isolates are clustered by phenotype and associated genomic variations are identified in large genome collections [50, 64, 100]. Early genome typing using DNA microarrays hinted at the level of diversity among *C. jejuni* isolates [27, 101], and comparisons of large isolate genome collections are now linking strain variation to differences in ecology [65, 102–105], epidemiology and evolution [63, 100, 106–110]. Advances in sequencing technology are helping us study genomes variation in greater depth and long read sequencing of isolate 2 identified large inversions (>90,000 bp) compared to the original finished genome (Table S1).

In conclusion, the genotypic and phenotypic differences among NCTC 11168 strains in this study, probably as a result of evolution during repeated passages, emphasises the need for laboratories to maintain isolate collections with detailed records and good culture practices. This essentially reaffirms the work of microbiology pioneers who developed practices to minimise variation between strains and laboratories. However, in the genomics era, it may also be prudent to sequence strains more routinely, particularly as the costs continue to decline. While the interpretation of experiments using reference type strains may be adapting to more detailed genomic data and improved understanding of genome evolution, the strains themselves remain an essential resource in microbiology. The perceived power of large-scale comparative genomics and statistical genetics studies typically lies in the ability to identify genes or genetic variation that confers putative functional differences to the bacterium. Confirming these associated gene functions [56] requires traditional microbiology based upon a detailed understanding of reliable reference type control strains such as NCTC 11168.

## Author statements

### Authors and contributors

Conceptualisation: SKS.

Formal analysis: BP, LKW, JKC, MDH, MD and JR.

Resources: SS, BSL, CC-U, EA, AV, CF, PE, DL, JAP, TAC, MPS, TSW, TJH, AJC, FMC, MCJM, KJF, NS, DJK, BMP, BWW, JP and AHMvV.

Data curation: BP, GM, KAJ, MCJM and SKS

Writing: BP, LKW and SKS. All authors contributed and approved the final manuscript.

### Ethical approval

Not applicable

### Consent for publication

Not applicable

## Acknowledgements

All high performance computing was performed on MRC CLIMB, funded by the Medical Research Council (MR/L015080/1). This publication made use of the PubMLST website (http://pubmlst.org/) developed by Keith Jolley and Martin Maiden (Jolley and Maiden, 2010) and sited at the University of Oxford. The development of that website was funded by the Wellcome Trust. We also thank all *Campylobacter* researchers who have maintained, cultured and disseminated this type strain since its deposition into the NCTC archives in 1977.

## Supplementary data

**Table S1:** Isolate list

**Table S2:** Isolates used for genomic context

**File S1:** Alignment file: NCTC11168 isolates and 83 previously published genomes.

**File S2:** Alignment file: NCTC11168 isolates only.

**File S3:** wgMLST

**File S4:** SNP matrix

## Data bibliography

1. Pascoe B, Williams LK, Calland JK, Méric G, Hitchings MD, Dyer M, Ryder J, Allen E, Vidal A, Fearnley C, Everest P, Linton D, Pachebat JA, Cogan TA, Stevens MP, Wilkinson TS, Humphrey TJ, Cody AJ, Colles FM, Jolley KA, Maiden MCJ, Forbes K, Strachan N, Kelly DJ, Pearson BM, Wren BW, Parkhill J, van Vliet AHM, Sheppard SK. Sequence Read Archive (SRA), BioProject Accession PRJNA517467 (2019).

2. Sheppard SK, Didelot X, Jolley KA, Darling AE, Pascoe B, Méric G, Kelly DJ, Cody AJ, Colles FM, Strachan NJ, Ogden ID, Forbes K, French NP, Carter P, Miller WG, McCarthy ND, Owen R, Litrup E, Egholm M, Affourtit JP, Bentley SD, Parkhill J, Maiden MCJ, Falush D. Sequence Read Archive (SRA), BioProject Accession PRJNA177352 (2013).

3. Sheppard SK, Didelot X, Méric G, Torralbo A, Jolley KA, Kelly DJ, Bentley SD, Maiden MCJ, Parkhill J, Falush D. European Nucleotide Archive (ENA), Study Accession ERP000129 (2013).

4. Pearson BM, Gaskin DJ, Segers RP, Wells JM, Nuijten PJ, van Vliet AH. GenBank sequence: CP000814.1 (2007).

5. Fouts D, Nelson K, Sebastian Y. GenBank sequence: CP000538.1 (2006).

6. Fouts DE, Mongodin EF, Mandrell RE, Miller WG, Rasko DA, Ravel J, Brinkac LM, DeBoy RT, Parker CT, Daugherty SC, Dodson RJ, Durkin AS, Madupu R, Sullivan SA, Shetty JU, Ayodeji MA, Shvartsbeyn A, Schatz MC, Badger JH, Fraser CM, Nelson KE. GenBank sequence: CP000025.1 (2004).

7. Gundogdu O, Bentley SD, Holden MT, Parkhill J, Dorrell N, Wren BW. GenBank sequence: AL111168.1 (2006).

## References

1. Méric G, Hitchings MD, Pascoe B, Sheppard SK. From Escherich to the Escherichia coli genome. Lancet Infect Dis 2016;16:634–636.

2. Skirrow MB. Campylobacter enteritis: a “new” disease. Br Med J 1977;2:9–11.

3. Kist M. [Who discovered Campylobacter jejuni/coli? A review of hitherto disregarded literature]. Zentralbl Bakteriol Mikrobiol Hyg A 1986;261:177–86.

4. Shulman ST, Friedmann HC, Sims RH. Theodor Escherich: The First Pediatric Infectious Diseases Physician? Clin Infect Dis 2007;45:1025–1029.

5. Dunne KA, Chaudhuri RR, Rossiter AE, Beriotto I, Browning DF, et al. Sequencing a piece of history: complete genome sequence of the original Escherichia coli strain. Microb Genomics;3. Epub ahead of print 23 March 2017. DOI: 10.1099/mgen.0.000106.

6. Altekruse SF, Swerdlow DL, Stern NJ. Microbial food borne pathogens. Campylobacter jejuni. Vet Clin North Am Food Anim Pract 1998;14:31–40.

7. Levy AJ. A gastro-enteritis cutbreak probably due to a bovine strain of vibrio. Yale J Biol Med 1946;18:243–58.

8. Jones FS, Orcutt M, Little RB. VIBRIOS (VIBRIO JEJUNI, N.SP.) ASSOCIATED WITH INTESTINAL DISORDERS OF COWS AND CALVES. J Exp Med 1931;53:853–63.

9. Smith T, Taylor MS. SOME MORPHOLOGICAL AND BIOLOGICAL CHARACTERS OF THE SPIRILLA (VIBRIO FETUS, N. SP.) ASSOCIATED WITH DISEASE OF THE FETAL MEMBRANES IN CATTLE. J Exp Med 1919;30:299–311.

10. Butzler J-P. Campylobacter, from obscurity to celebrity. Clin Microbiol Infect 2004;10:868–876.

11. Dekeyser P, Gossuin-Detrain M, Butzler JP, Sternon J. Acute enteritis due to related vibrio: first positive stool cultures. J Infect Dis 1972;125:390–2.

12. Butzler JP, Dekeyser P, Detrain M, Dehaen F. Related vibrio in stools. J Pediatr 1973;82:493–5.

13. Cadranel S, Rodesch P, Butzler JP, Dekeyser P. Enteritis due to “related vibrio” in children. Am J Dis Child 1973;126:152–5.

14. Sebald M, Veron M. [BASE DNA CONTENT AND CLASSIFICATION OF VIBRIOS]. Ann Inst Pasteur (Paris) 1963;105:897–910.

15. Veron M, Chatelain R. Taxonomic Study of the Genus Campylobacter Sebald and Veron and Designation of the Neotype Strain for the Type Species, Campylobacter fetus (Smith and Taylor) Sebald and Veron. Int J Syst Bacteriol 1973;23:122–134.

16. Skirrow MB, Benjamin J. Differentiation of enteropathogenic Campylobacter. J Clin Pathol 1980;33:1122.

17. Skirrow MB, Benjamin J. ‘1001’ Campylobacters: cultural characteristics of intestinal campylobacters from man and animals. J Hyg (Lond) 1980;85:427–42.

18. Woodward DL, Rodgers FG. Identification of Campylobacter heat-stable and heat-labile antigens by combining the Penner and Lior serotyping schemes. J Clin Microbiol 2002;40:741–5.

19. Taylor DE, Chang N. Immunoblot and enzyme-linked immunosorbent assays of Campylobacter major outer-membrane protein and application to the differentiation of Campylobacter species. Mol Cell Probes 1987;1:261–74.

20. Lior H. New, extended biotyping scheme for Campylobacter jejuni, Campylobacter coli, and “Campylobacter laridis”. J Clin Microbiol 1984;20:636–40.

21. Taylor DE, Eaton M, Yan W, Chang N. Genome maps of Campylobacter jejuni and Campylobacter coli. J Bacteriol 1992;174:2332–7.

22. Newnham E, Chang N, Taylor DE. Expanded genomic map of *Campylobacter jejuni* UA580 and localization of 23S ribosomal rRNA genes by I- *CeuI* restriction endonuclease digestion. FEMS Microbiol Lett 1996;142:223–229.

23. Karlyshev A V., Henderson J, Ketley JM, Wren BW. An improved physical and genetic map of Campylobacter jejuni NCTC 11168 (UA580). Microbiology 1998;144:503–508.

24. Parkhill J, Wren BW, Mungall K, Ketley JM, Churcher C, et al. The genome sequence of the food-borne pathogen Campylobacter jejuni reveals hypervariable sequences. Nature 2000;403:665–668.

25. Revez J, Schott T, Rossi M, Hänninen M-L. Complete genome sequence of a variant of Campylobacter jejuni NCTC 11168. J Bacteriol 2012;194:6298–9.

26. Gundogdu O, Bentley SD, Holden MT, Parkhill J, Dorrell N, et al. Re-annotation and re-analysis of the Campylobacter jejuni NCTC11168 genome sequence. BMC Genomics 2007;8:162.

27. Dorrell N, Mangan JA, Laing KG, Hinds J, Linton D, et al. Whole Genome Comparison of Campylobacter jejuni Human Isolates Using a Low-Cost Microarray Reveals Extensive Genetic Diversity. Genome Res 2001;11:1706–1715.

28. Gaynor EC, Cawthraw S, Manning G, MacKichan JK, Falkow S, et al. The genome-sequenced variant of Campylobacter jejuni NCTC 11168 and the original clonal clinical isolate differ markedly in colonization, gene expression, and virulence-associated phenotypes. J Bacteriol 2004;186:503–17.

29. Cooper KK, Cooper MA, Zuccolo A, Joens LA. Re-sequencing of a virulent strain of Campylobacter jejuni NCTC11168 reveals potential virulence factors. Res Microbiol 2013;164:6–11.

30. Szymanski CM, Logan SM, Linton D, Wren BW. Campylobacter--a tale of two protein glycosylation systems. Trends Microbiol 2003;11:233–8.

31. Thibault P, Logan SM, Kelly JF, Brisson J-R, Ewing CP, et al. Identification of the Carbohydrate Moieties and Glycosylation Motifs in *Campylobacter jejuni* Flagellin. J Biol Chem 2001;276:34862–34870.

32. Linton D, Karlyshev A V., Hitchen PG, Morris HR, Dell A, et al. Multiple N-acetyl neuraminic acid synthetase (neuB) genes in Campylobacter jejuni: identification and characterization of the gene involved in sialylation of lipo-oligosaccharide. Mol Microbiol 2000;35:1120–1134.

33. Guerry P, Ewing CP, Hickey TE, Prendergast MM, Moran AP. Sialylation of lipooligosaccharide cores affects immunogenicity and serum resistance of Campylobacter jejuni. Infect Immun 2000;68:6656–62.

34. Howard SL, Jagannathan A, Soo EC, Hui JPM, Aubry AJ, et al. Campylobacter jejuni glycosylation island important in cell charge, legionaminic acid biosynthesis, and colonization of chickens. Infect Immun 2009;77:2544–56.

35. Linton D, Karlyshev A V, Wren BW. Deciphering Campylobacter jejuni cell surface interactions from the genome sequence. Curr Opin Microbiol 2001;4:35–40.

36. Jones MA, Marston KL, Woodall CA, Maskell DJ, Linton D, et al. Adaptation of Campylobacter jejuni NCTC11168 to high-level colonization of the avian gastrointestinal tract. Infect Immun 2004;72:3769–76.

37. Carrillo CD, Taboada E, Nash JHE, Lanthier P, Kelly J, et al. Genome-wide Expression Analyses of *Campylobacter jejuni* NCTC11168 Reveals Coordinate Regulation of Motility and Virulence by *flhA*. J Biol Chem 2004;279:20327–20338.

38. Perera VN, Nachamkin I, Ung H, Patterson JH, McConville MJ, et al. Molecular mimicry in *Campylobacter jejuni*: role of the lipo-oligosaccharide core oligosaccharide in inducing anti-ganglioside antibodies. FEMS Immunol Med Microbiol 2007;50:27–36.

39. Holmes K, Mulholland F, Pearson BM, Pin C, McNicholl-Kennedy J, et al. Campylobacter jejuni gene expression in response to iron limitation and the role of Fur. Microbiology 2005;151:243–257.

40. Palyada K, Threadgill D, Stintzi A. Iron acquisition and regulation in Campylobacter jejuni. J Bacteriol 2004;186:4714–29.

41. Xu F, Zeng X, Haigh RD, Ketley JM, Lin J. Identification and characterization of a new ferric enterobactin receptor, CfrB, in Campylobacter. J Bacteriol 2010;192:4425– 35.

42. Nielsen LN, Luijkx TA, Vegge CS, Johnsen CK, Nuijten P, et al. Identification of immunogenic and virulence-associated Campylobacter jejuni proteins. Clin Vaccine Immunol 2012;19:113–9.

43. Mandal RK, Jiang T, Kwon YM. Essential genome of Campylobacter jejuni. BMC Genomics 2017;18:616.

44. de Vries SP, Gupta S, Baig A, Wright E, Wedley A, et al. Genome-wide fitness analyses of the foodborne pathogen Campylobacter jejuni in in vitro and in vivo models. Sci Rep 2017;7:1251.

45. Wright JA, Grant AJ, Hurd D, Harrison M, Guccione EJ, et al. Metabolite and transcriptome analysis of Campylobacter jejuni in vitro growth reveals a stationary-phase physiological switch. Microbiology 2009;155:80–94.

46. Guccione EJ, Kendall JJ, Hitchcock A, Garg N, White MA, et al. Transcriptome and proteome dynamics in chemostat culture reveal how *Campylobacter jejuni* modulates metabolism, stress responses and virulence factors upon changes in oxygen availability. Environ Microbiol 2017;19:4326–4348.

47. Kalmokoff M, Lanthier P, Tremblay T-L, Foss M, Lau PC, et al. Proteomic analysis of Campylobacter jejuni 11168 biofilms reveals a role for the motility complex in biofilm formation. J Bacteriol 2006;188:4312–20.

48. Reuter M, Mallett A, Pearson BM, Vliet AHM van. Biofilm Formation by Campylobacter jejuni Is Increased under Aerobic Conditions. Appl Environ Microbiol 2010;76:2122–2128.

49. Brown HL, Hanman K, Reuter M, Betts RP, van Vliet AHM. Campylobacter jejuni biofilms contain extracellular DNA and are sensitive to DNase I treatment. Front Microbiol 2015;6:699.

50. Pascoe B, Méric G, Murray S, Yahara K, Mageiros L, et al. Enhanced biofilm formation and multi-host transmission evolve from divergent genetic backgrounds in Campylobacter jejuni. Environ Microbiol 2015;17:4779–4789.

51. Oh E, Andrews KJ, Jeon B. Enhanced Biofilm Formation by Ferrous and Ferric Iron Through Oxidative Stress in Campylobacter jejuni. Front Microbiol 2018;9:1204.

52. Karlyshev A V, Linton D, Gregson NA, Lastovica AJ, Wren BW. Genetic and biochemical evidence of a Campylobacter jejuni capsular polysaccharide that accounts for Penner serotype specificity. Mol Microbiol 2000;35:529–41.

53. Atack JM, Kelly DJ. Oxidative stress in *Campylobacter jejuni* : responses, resistance and regulation. Future Microbiol 2009;4:677–690.

54. Kendall JJ, Barrero-Tobon AM, Hendrixson DR, Kelly DJ. Hemerythrins in the microaerophilic bacterium *C ampylobacter jejuni* help protect key iron-sulphur cluster enzymes from oxidative damage. Environ Microbiol 2014;16:1105–1121.

55. Thomas DK, Lone AG, Selinger LB, Taboada EN, Uwiera RRE, et al. Comparative Variation within the Genome of Campylobacter jejuni NCTC 11168 in Human and Murine Hosts. PLoS One 2014;9:e88229.

56. Yahara K, Méric G, Taylor AJ, de Vries SPW, Murray S, et al. Genome-wide association of functional traits linked with Campylobacter jejuni survival from farm to fork. Environ Microbiol;19. Epub ahead of print 2017. DOI: 10.1111/1462-2920.13628.

57. Sheppard SK, Guttman DS, Fitzgerald JR. Population genomics of bacterial host adaptation. Nat Rev Genet 2018;19:549–565.

58. Sleight SC, Orlic C, Schneider D, Lenski RE. Genetic Basis of Evolutionary Adaptation by *Escherichia coli* to Stressful Cycles of Freezing, Thawing and Growth. Genetics 2008;180:431–443.

59. Sproston EL, Wimalarathna HML, Sheppard SK. Trends in fluoroquinolone resistance in Campylobacter. Microb Genomics;4. Epub ahead of print 1 August 2018. DOI: 10.1099/mgen.0.000198.

60. Viana D, Comos M, McAdam PR, Ward MJ, Selva L, et al. A single natural nucleotide mutation alters bacterial pathogen host tropism. Nat Genet 2015;47:361– 366.

61. Jolley KA, Maiden MCJ. BIGSdb: Scalable analysis of bacterial genome variation at the population level. BMC Bioinformatics 2010;11:595.

62. Sheppard SK, Jolley KA, Maiden MCJ. A Gene-By-Gene Approach to Bacterial Population Genomics: Whole Genome MLST of Campylobacter. Genes (Basel) 2012;3:261–277.

63. Sheppard SK, Didelot X, Jolley KA, Darling AE, Pascoe B, et al. Progressive genome-wide introgression in agricultural *Campylobacter coli*. Mol Ecol 2013;22:1051–1064.

64. Sheppard SK, Didelot X, Meric G, Torralbo A, Jolley KA, et al. Genome-wide association study identifies vitamin B5 biosynthesis as a host specificity factor in Campylobacter. Proc Natl Acad Sci U S A 2013;110:11923–11927.

65. Sheppard SK, Cheng L, Méric G, De Haan CPA, Llarena AK, et al. Cryptic ecology among host generalist Campylobacter jejuni in domestic animals. Mol Ecol. Epub ahead of print 2014. DOI: 10.1111/mec.12742.

66. Méric G, Yahara K, Mageiros L, Pascoe B, Maiden MCJ, et al. A reference pan-genome approach to comparative bacterial genomics: Identification of novel epidemiological markers in pathogenic Campylobacter. PLoS One;9. Epub ahead of print 2014. DOI: 10.1371/journal.pone.0092798.

67. Altschul SF, Gish W, Miller W, Myers EW, Lipman DJ. Basic local alignment search tool. J Mol Biol 1990;215:403–410.

68. Katoh K, Standley DM. MAFFT Multiple Sequence Alignment Software Version 7: Improvements in Performance and Usability. Mol Biol Evol 2013;30:772–780.

69. Price MN, Dehal PS, Arkin AP. FastTree 2 - Approximately maximum-likelihood trees for large alignments. PLoS One;5. Epub ahead of print 2010. DOI: 10.1371/journal.pone.0009490.

70. Didelot X, Wilson DJ. ClonalFrameML: Efficient Inference of Recombination in Whole Bacterial Genomes. PLoS Comput Biol;11. Epub ahead of print 2015. DOI: 10.1371/journal.pcbi.1004041.

71. Croucher NJ, Page AJ, Connor TR, Delaney AJ, Keane JA, et al. Rapid phylogenetic analysis of large samples of recombinant bacterial whole genome sequences using Gubbins. Nucleic Acids Res 2015;43:e15–e15.

72. Seemann T. Snippy: Fast bacterial variant calling from NGS reads. https://github.com/tseemann/snippy (2015).

73. Seemann T. Prokka: rapid prokaryotic genome annotation. Bioinformatics 2014;30:2068–2069.

74. Grüning B, Dale R, Sjödin A, Chapman BA, Rowe J, et al. Bioconda: sustainable and comprehensive software distribution for the life sciences. Nat Methods 2018;15:475–476.

75. Connor TR, Loman NJ, Thompson S, Smith A, Southgate J, et al. CLIMB (the Cloud Infrastructure for Microbial Bioinformatics): an online resource for the medical microbiology community. Microb Genomics;2. Epub ahead of print 20 September 2016. DOI: 10.1099/mgen.0.000086.

76. Welch BL. On the Comparison of Several Mean Values: An Alternative Approach. Biometrika 1951;38:330.

77. Guerry P, Doig P, Alm RA, Burr DH, Kinsella N, et al. Identification and characterization of genes required for post-translational modification of Campylobacter coli VC167 flagellin. Mol Microbiol 1996;19:369–78.

78. Golden NJ, Acheson DWK. Identification of motility and autoagglutination Campylobacter jejuni mutants by random transposon mutagenesis. Infect Immun 2002;70:1761–71.

79. Culebro A, Revez J, Pascoe B, Friedmann Y, Hitchings MD, et al. Large Sequence Diversity within the Biosynthesis Locus and Common Biochemical Features of Campylobacter coli Lipooligosaccharides. J Bacteriol 2016;198:2829–40.

80. Guerry P, Ewing CP, Schirm M, Lorenzo M, Kelly J, et al. Changes in flagellin glycosylation affect Campylobacter autoagglutination and virulence. Mol Microbiol 2006;60:299–311.

81. Wu Z, Periaswamy B, Sahin O, Yaeger M, Plummer P, et al. Point mutations in the major outer membrane protein drive hypervirulence of a rapidly expanding clone of *Campylobacter jejuni*. Proc Natl Acad Sci 2016;113:10690–10695.

82. Parker CT, Gilbert M, Yuki N, Endtz HP, Mandrell RE. Characterization of lipooligosaccharide-biosynthetic loci of Campylobacter jejuni reveals new lipooligosaccharide classes: evidence of mosaic organizations. J Bacteriol 2008;190:5681–9.

83. Backert S, Hofreuter D. Molecular methods to investigate adhesion, transmigration, invasion and intracellular survival of the foodborne pathogen Campylobacter jejuni. J Microbiol Methods 2013;95:8–23.

84. Karlyshev A V., Linton D, Gregson NA, Wren BW. A novel paralogous gene family involved in phase-variable flagella-mediated motility in Campylobacter jejuni. Microbiology 2002;148:473–480.

85. Hendrixson DR, DiRita VJ. Transcription of sigma54-dependent but not sigma28- dependent flagellar genes in Campylobacter jejuni is associated with formation of the flagellar secretory apparatus. Mol Microbiol 2003;50:687–702.

86. Novik V, Hofreuter D, Galan JE. Identification of Campylobacter jejuni Genes Involved in Its Interaction with Epithelial Cells. Infect Immun 2010;78:3540–3553.

87. Reid AN, Pandey R, Palyada K, Naikare H, Stintzi A. Identification of Campylobacter jejuni Genes Involved in the Response to Acidic pH and Stomach Transit. Appl Environ Microbiol 2008;74:1583–1597.

88. Hitchen P, Brzostek J, Panico M, Butler JA, Morris HR, et al. Modification of the Campylobacter jejuni flagellin glycan by the product of the Cj1295 homopolymerictract-containing gene. Microbiology 2010;156:1953–1962.

89. Aidley J, Rajopadhye S, Akinyemi NM, Lango-Scholey L, Bayliss CD, et al. Nonselective Bottlenecks Control the Divergence and Diversification of Phase-Variable Bacterial Populations. MBio;8. Epub ahead of print 3 May 2017. DOI: 10.1128/mBio.02311-16.

90. Aidley J, Wanford JJ, Green LR, Sheppard SK, Bayliss CD. PhasomeIt: an ‘omics’ approach to cataloguing the potential breadth of phase variation in the genus Campylobacter. Microb Genomics;4. Epub ahead of print 1 November 2018. DOI: 10.1099/mgen.0.000228.

91. Kearns DB. A field guide to bacterial swarming motility. Nat Rev Microbiol 2010;8:634–44.

92. Adler J, Templeton B. The Effect of Environmental Conditions on the Motility of Escherichia coli. J Gen Microbiol 1967;46:175–184.

93. Navarro Llorens JM, Tormo A, Martínez-García E. Stationary phase in gram-negative bacteria. FEMS Microbiol Rev 2010;34:476–495.

94. Rendueles O, Velicer GJ. Evolution by flight and fight: diverse mechanisms of adaptation by actively motile microbes. ISME J 2016;11:555–568.

95. Konkel ME, Klena JD, Rivera-Amill V, Monteville MR, Biswas D, et al. Secretion of Virulence Proteins from Campylobacter jejuni Is Dependent on a Functional Flagellar Export Apparatus. J Bacteriol 2004;186:3296–3303.

96. Guerry P. Campylobacter flagella: not just for motility. Trends Microbiol 2007;15:456–461.

97. Revez J, Schott T, Llarena A-K, Rossi M, Hänninen M-L. Genetic heterogeneity of Campylobacter jejuni NCTC 11168 upon human infection. Infect Genet Evol 2013;16:305–9.

98. Clinical and Laboratory Standards. Performance Standards for Antimicrobial Susceptibility Testing An informational supplement for global application developed through the Clinical and Laboratory Standards Institute. CLSI Doc M100-S16CLSI, Wayne, PA. www.clsi.org. (2006, accessed 19 March 2019).

99. Draper JL, Hansen LM, Bernick DL, Abedrabbo S, Underwood JG, et al. Fallacy of the Unique Genome: Sequence Diversity within Single *Helicobacter pylori* Strains. MBio;8. Epub ahead of print 8 March 2017. DOI: 10.1128/mBio.02321-16.

100. Pascoe B, Méric G, Yahara K, Wimalarathna H, Murray S, et al. Local genes for local bacteria: Evidence of allopatry in the genomes of transatlantic *Campylobacter* populations. Mol Ecol 2017;26:4497–4508.

101. Champion OL, Gaunt MW, Gundogdu O, Elmi A, Witney AA, et al. Comparative phylogenomics of the food-borne pathogen Campylobacter jejuni reveals genetic markers predictive of infection source. Proc Natl Acad Sci U S A 2005;102:16043–8.

102. Sheppard SK, Dallas JF, Wilson DJ, Strachan NJC, McCarthy ND, et al. Evolution of an Agriculture-Associated Disease Causing Campylobacter coli Clade: Evidence from National Surveillance Data in Scotland. PLoS One 2010;5:e15708.

103. Sheppard SK, Colles FM, McCarthy ND, Strachan NJC, Ogden ID, et al. Niche segregation and genetic structure of Campylobacter jejuni populations from wild and agricultural host species. Mol Ecol 2011;20:3484–3490.

104. Méric G, McNally A, Pessia A, Mourkas E, Pascoe B, et al. Convergent Amino Acid Signatures in Polyphyletic Campylobacter jejuni Subpopulations Suggest Human Niche Tropism. Genome Biol Evol 2018;10:763–774.

105. Woodcock DJ, Krusche P, Strachan NJC, Forbes KJ, Cohan FM, et al. Genomic plasticity and rapid host switching can promote the evolution of generalism: A case study in the zoonotic pathogen Campylobacter. Sci Rep. Epub ahead of print 2017. DOI: 10.1038/s41598-017-09483-9.

106. Sheppard SK, McCarthy ND, Falush D, Maiden MCJ. Convergence of Campylobacter Species: Implications for Bacterial Evolution. Science (80-) 2008;320:237–239.

107. Sheppard SK, McCarthy ND, Jolley KA, Maiden MCJ. Introgression in the genus Campylobacter: generation and spread of mosaic alleles. Microbiology 2011;157:1066–1074.

108. Dearlove BL, Cody AJ, Pascoe B, Méric G, Wilson DJ, et al. Rapid host switching in generalist Campylobacter strains erodes the signal for tracing human infections. ISME J;10. Epub ahead of print 2016. DOI: 10.1038/ismej.2015.149.

109. Baily JL, Méric G, Bayliss S, Foster G, Moss SE, et al. Evidence of land-sea transfer of the zoonotic pathogen Campylobacter to a wildlife marine sentinel species. Mol Ecol;24. Epub ahead of print 2015. DOI: 10.1111/mec.13001.

110. Thépault A, Méric G, Rivoal K, Pascoe B, Mageiros L, et al. Genome-wide identification of host-segregating epidemiological markers for source attribution in Campylobacter jejuni. Appl Environ Microbiol;83. Epub ahead of print 2017. DOI: 10.1128/AEM.03085-16.

