## Supplemental Table 1 for "Domestication of *Campylobacter jejuni* NCTC 11168"

| Isolate | ID | Source laboratory | Variant / comment | Original source | Archived | Number of contigs | Genome size (bp) | N50 contig length (L50) | GC% | SRA accession | BioProject |
|---|---|---|---|---|---|---|---|---|---|---|---|
| 1 | 5920 | Aberystwyth | Primary lab strain | NCTC | 2015 | 18 | 1,626,801 | 174,210 | 30.5 | SRR8731760 | PRJNA517467 |
| 2 | 5921 | Aberdeen | Primary lab strain | NCTC | 2002 | 16 | 1,626,067 | 174,212 | 30.5 | SRR8731759 | PRJNA517467 |
| 3 | 5922 | Bristol | Non-motile | NCTC | 2002 | 36 | 1,634,599 | 189,487 | 30.5 | SRR8731758 | PRJNA517467 |
| 4 | 5923 | Bristol | Hyper-motile | NCTC | 2005 | 18 | 1,626,519 | 154,057 | 30.5 | SRR8731757 | PRJNA517467 |
| 5 | 5925 | Glasgow | Hyper-motile | London | 2000 | 22 | 1,625,874 | 173,183 | 30.5 | SRR8731764 | PRJNA517467 |
| 6 | 5926 | Glasgow | Original strain | NCTC (via Martin Skirrow) | 2000 | 16 | 1,625,293 | 154,057 | 30.5 | SRR8731763 | PRJNA517467 |
| 7 | 5927 | Glasgow | Sequenced variant | London | 2000 | 23 | 1,626,367 | 173,039 | 30.5 | SRR8731762 | PRJNA517467 |
| 8 | 5928 | Norwich | Primary lab strain | NCTC | 2004 | 19 | 1,626,763 | 188,103 | 30.5 | SRR8731761 | PRJNA517467 |
| 9 | 5929 | Norwich | Hyper-motile | NCTC | 2004 | 31 | 1,625,378 | 100,444 | 30.5 | SRR8731766 | PRJNA517467 |
| 10 | 5930 | London | -- | London | 2000 | 35 | 1,624,738 | 108,757 | 30.5 | SRR8731765 | PRJNA517467 |
| 11 | 5931 | London | Hyper-motile | London | 2000 | 20 | 1,641,300 | 188,163 | 30.5 | SRR8731768 | PRJNA517467 |
| 12 | 5932 | Manchester | Hyper-motile | London | 2003 | 25 | 1,628,343 | 188,003 | 30.5 | SRR8731767 | PRJNA517467 |
| 13 | 5933 | Swansea | Recently purchased | NCTC | 2016 | 16 | 1,694,909 | 153,963 | 30.5 | tbc | tbc |
| 14 | 5934 | Oxford | Primary lab strain | NCTC | 2014 | 19 | 1,625,944 | 122,646 | 30.5 | SRR8731770 | PRJNA517467 |
| 15 | 5935 | Sheffield | Primary lab strain | London | 2015 | 14 | 1,625,814 | 189,487 | 30.5 | SRR8731769 | PRJNA517467 |
| 16 | 5936 | Sheffield | Hyper-motile | London | 2013 | 14 | 1,626,210 | 189,487 | 30.5 | SRR8731772 | PRJNA517467 |
| 17 | 5937 | Sheffield | WT-2000 | London (via Birmingham) | 2000 | 15 | 1,612,402 | 189,478 | 30.5 | SRR8731771 | PRJNA517467 |
| 18 | 5938 | Sheffield | WT-2010 (subcultured from WT-2000) | London | 2010 | 13 | 1,625,308 | 189,488 | 30.5 | SRR8731774 | PRJNA517467 |
| 19 | 5939 | London | Hyper-motile | London | 2002 | 14 | 1,625,123 | 189,490 | 30.5 | SRR8731773 | PRJNA517467 |
| 20 | 5940 | London | [Genome previously sequenced] | London | 2002 | 18 | 1,625,478 | 189,490 | 30.5 | tbc | tbc |
| 21 | 5941 | Surrey | Primary lab strain | NCTC (via Cambridge) | 2000 | 16 | 1,625,755 | 271,714 | 30.5 | SRR8731776 | PRJNA517467 |
| 22 | 5942 | Surrey | -- | NCTC (via Cambridge) | 2000 | 16 | 1,624,913 | 154,057 | 30.5 | SRR8731775 | PRJNA517467 |
| 23 | 5943 | Edinburgh | -- | London (via Sheffield) | 2013 | 20 | 1,626,490 | 189,488 | 30.5 | SRR8731777 | PRJNA517467 |
| Reference | -- | NCTC | Original sequenced isolate | London | 2000 | 1 | 1,641,481 | 1,641,481 | 30.5 | EMBL: AL11168.1 | -- |
