## Supplemental Table 2 for "Domestication of *Campylobacter jejuni* NCTC 11168"

| ID | Isolate | Country | Year | Source | Genome size (bp) | Contigs | ST | Clonal complex | BioProject | Accession |
|---|---|---|---|---|---|---|---|---|---|---|
| 4 | CAMP45 | UK | 2005 | chicken | 1,596,969 | 108 | 45 | ST-45 | PRJNA177352 | ANGO00000000.1 |
| 22 | CAMP2488 | UK | 2001 | chicken | 1,573,736 | 169 | 257 | ST-257 | ERP000129 | ERS007807 |
| 26 | NCTC11828 | UK | 2007 | Lab strain | 1,628,115 | 1 | 267 | ST-283 | PRJNA17953 | CP000814.1 |
| 27 | NC 008787 | USA | 2007 | Lab strain | 1,616,554 | 1 | 604 | ST-42 | PRJNA224116 | CP000538.1 |
| 28 | RM1221 | USA | 2002 | Lab strain | 1,777,831 | 1 | 354 | ST-354 | PRJNA224116 | CP000025.1 |
| **29** | **NCTC11168** | **UK** | **2000** | Lab strain | **1,641,481** | **1** | **43** | **ST-21** | **PRJNA57587** | **AL111168.1** |
| 30 | CAMP1044 | UK | 2007 | Lab strain | 1,613,621 | 465 | -- | -- | PRJNA177352 | ANHK00000000.1 |
| 32 | CampsClin11 | UK | 2005 | clinical | 1,650,105 | 92 | 11 | ST-45 | ERP000129 | ERR024478 |
| 34 | CampsClin262 | UK | 2005 | clinical | 1,643,032 | 91 | 262 | ST-21 | ERP000129 | ERR024475 |
| 36 | CampsClin266 | UK | 2006 | clinical | 1,695,272 | 47 | 266 | ST-21 | ERP000129 | ERR024476 |
| 37 | CampsClin883 | UK | 2006 | clinical | 1,667,560 | 113 | 883 | ST-21 | ERP000129 | ERR024477 |
| 39 | chick2219 | UK | 2005 | chicken | 1,616,482 | 84 | 2219 | ST-45 | ERP000129 | ERR024426 |
| 40 | chicka21 | UK | 2006 | chicken | 1,726,327 | 195 | 21 | ST-21 | ERP000129 | ERR024431 |
| 42 | cow42 | UK | 2006 | cattle | 1,672,737 | 41 | 42 | ST-42 | ERP000129 | ERR024430 |
| 45 | chick594 | UK | 2006 | chicken | 1,609,163 | 72 | 583 | ST-45 | ERP000129 | ERR024434 |
| 47 | cow2674 | UK | 2006 | cattle | 3,453,740 | 455 | 21 | ST-21 | ERP000129 | ERR024436 |
| 48 | cow206 | UK | 2006 | cattle | 1,671,619 | 62 | 206 | ST-206 | ERP000129 | ERR024437 |
| 49 | cow38 | UK | 2006 | cattle | 1,663,148 | 181 | 38 | ST-48 | ERP000129 | ERR024427 |
| 52 | cow334 | UK | 2006 | cattle | 1,616,567 | 102 | 334 | ST-45 | ERP000129 | ERR023263 |
| 54 | chick267 | UK | 2005 | chicken | 1,591,217 | 241 | 267 | ST-283 | ERP000129 | ERR023268 |
| 55 | CampsClin230 | UK | 2006 | clinical | 1,625,711 | 271 | 230 | ST-45 | ERP000129 | ERR024480 |
| 56 | cowa45 | UK | 2006 | cattle | 1,607,778 | 62 | 45 | ST-45 | ERP000129 | ERR023269 |
| 57 | chick2213 | UK | 2005 | chicken | 1,620,325 | 158 | 334 | ST-45 | ERP000129 | ERR023270 |
| 59 | cow518 | UK | 2006 | cattle | 1,705,325 | 60 | 21 | ST-21 | ERP000129 | ERR024441 |
| 60 | CampsClin53 | UK | 2005 | clinical | 1,658,292 | 58 | 53 | ST-21 | ERP000129 | ERR024474 |
| 62 | cowa21 | UK | 2006 | cattle | 1,658,436 | 100 | 21 | ST-21 | ERP000129 | ERR023273 |
| 63 | chickc21 | UK | 2006 | chicken | 1,679,349 | 139 | 21 | ST-21 | ERP000129 | ERR024447 |
| 64 | chick25 | UK | 2006 | chicken | 1,698,035 | 113 | 814 | ST-661 | ERP000129 | ERR024448 |
| 65 | chick104 | UK | 2006 | chicken | 1,761,202 | 112 | 104 | ST-21 | ERP000129 | ERR023274 |
| 66 | chick353 | UK | 2009 | chicken | 1,776,210 | 129 | 353 | ST-353 | ERP000129 | ERR023264 |
| 67 | chickb354 | UK | 2009 | chicken | 1,688,706 | 143 | 354 | ST-354 | ERP000129 | ERR024453 |
| 68 | chick573 | UK | 2009 | chicken | 1,838,022 | 251 | 573 | ST-573 | ERP000129 | ERR023265 |
| 69 | chick2568 | UK | 2009 | chicken | 1,821,236 | 144 | 2568 | ST-661 | ERP000129 | ERR024456 |
| 70 | chickc45 | UK | 2009 | chicken | 1,595,762 | 343 | 45 | ST-45 | ERP000129 | ERR023266 |
| 71 | chick19 | UK | 2009 | chicken | 1,689,713 | 84 | 50 | ST-21 | ERP000129 | ERR023276 |
| 72 | chick50 | UK | 2009 | chicken | 1,692,341 | 63 | 50 | ST-21 | ERP000129 | ERR023280 |
| 73 | chick53 | UK | 2009 | chicken | 1,651,079 | 102 | 53 | ST-21 | ERP000129 | ERR023281 |
| 74 | chick262 | UK | 2009 | chicken | 1,606,379 | 65 | 262 | ST-21 | ERP000129 | ERR023282 |
| 75 | chick266 | UK | 2009 | chicken | 1,693,845 | 76 | 266 | ST-21 | ERP000129 | ERR023283 |
| 77 | chick1086 | UK | 2009 | chicken | 1,692,435 | 65 | 50 | ST-21 | ERP000129 | ERR023285 |
| 78 | chick1360 | UK | 2009 | chicken | 1,693,941 | 67 | 50 | ST-21 | ERP000129 | ERR023286 |
| 79 | chick11 | UK | 2009 | chicken | 1,645,238 | 120 | 11 | ST-45 | ERP000129 | ERR023287 |
| 80 | chick137 | UK | 2009 | chicken | 1,734,017 | 73 | 2030 | ST-257 | ERP000129 | ERR023277 |
| 81 | chick1003 | UK | 2009 | chicken | 1,617,200 | 106 | 1003 | ST-45 | ERP000129 | ERR023278 |
| 82 | chick2048 | UK | 2009 | chicken | 1,631,119 | 163 | 45 | ST-45 | ERP000129 | ERR024450 |
| 84 | chick2223 | UK | 2009 | chicken | 1,605,483 | 58 | 45 | ST-45 | ERP000129 | ERR024452 |
| 85 | cow3583 | UK | 2003 | cattle | 1,654,563 | 93 | 3583 | ST-42 | ERP000129 | ERR024442 |
| 87 | cow273 | UK | 2003 | cattle | 1,715,399 | 62 | 273 | ST-206 | ERP000129 | ERR024445 |
| 88 | cow270 | UK | 2003 | cattle | 1,716,038 | 72 | 270 | ST-403 | ERP000129 | ERR023293 |
| 89 | cowb21 | UK | 2003 | cattle | 1,659,711 | 130 | 21 | ST-21 | ERP000129 | ERR023294 |
| 90 | cowb45 | UK | 2003 | cattle | 1,603,131 | 64 | 45 | ST-45 | ERP000129 | ERR023295 |
| 91 | cowc45 | UK | 2003 | cattle | 1,602,224 | 47 | 45 | ST-45 | ERP000129 | ERR024443 |
| 92 | cowd45 | UK | 2003 | cattle | 1,607,691 | 98 | 45 | ST-45 | ERP000129 | ERR024444 |
| 94 | cow104 | UK | 2003 | cattle | 1,762,939 | 82 | 104 | ST-21 | ERP000129 | ERR024439 |
| 97 | cow3201 | UK | 2003 | cattle | 1,629,692 | 114 | 19 | ST-21 | ERP000129 | ERR023299 |
| 99 | cow3205 | UK | 2003 | cattle | 1,720,953 | 64 | 206 | ST-206 | ERP000129 | ERR023290 |
| 100 | cow137 | UK | 2003 | cattle | 1,626,334 | 192 | 137 | ST-45 | ERP000129 | ERR023291 |
| 102 | cow583 | UK | 2003 | cattle | 1,607,515 | 63 | 583 | ST-45 | ERP000129 | ERR027216 |
| 103 | cow3207 | UK | 2003 | cattle | 1,641,790 | 70 | 334 | ST-45 | ERP000129 | ERR027220 |
| 104 | cow3214 | UK | 2003 | cattle | 1,654,338 | 115 | 45 | ST-45 | ERP000129 | ERR027221 |
| 105 | chick354 | UK | 2004 | chicken | 1,697,013 | 75 | 257 | ST-257 | ERP000129 | ERR027222 |
| 106 | chick51 | UK | 2005 | chicken | 1,714,044 | 58 | 51 | ST-443 | ERP000129 | ERR024454 |
| 107 | chick1079 | UK | 2004 | chicken | 1,838,061 | 273 | 1079 | ST-573 | ERP000129 | ERR024455 |
| 108 | chick574 | UK | 2004 | chicken | 1,743,461 | 80 | 574 | ST-574 | ERP000129 | ERR027223 |
| 109 | chick814 | UK | 2004 | chicken | 1,759,789 | 152 | 814 | ST-661 | ERP000129 | ERR027224 |
| 110 | chickb21 | UK | 2003 | chicken | 1,656,261 | 72 | 21 | ST-21 | ERP000129 | ERR027225 |
| 111 | chickb45 | UK | 2004 | chicken | 1,649,834 | 80 | 45 | ST-45 | ERP000129 | ERR027226 |
| 112 | chickd45 | UK | 2004 | chicken | 1,618,948 | 55 | 45 | ST-45 | ERP000129 | ERR024440 |
| 113 | chick883 | UK | 2004 | chicken | 1,665,144 | 72 | 883 | ST-21 | ERP000129 | ERR027227 |
| 114 | chick230 | UK | 2004 | chicken | 1,633,592 | 79 | 230 | ST-45 | ERP000129 | ERR027217 |
| 116 | CampsClin21 | UK | 2005 | clinical | 1,656,471 | 65 | 9092 | ST-21 | ERP000129 | ERR024472 |
| 117 | OxClina21 | UK | 2003 | clinical | 1,697,696 | 113 | 21 | ST-21 | ERP000129 | ERR024457 |
| 119 | OxClina45 | UK | 2003 | clinical | 1,621,668 | 66 | 45 | ST-45 | ERP000129 | ERR024459 |
| 122 | starling177 | UK | 2007 | starling | 1,582,720 | 52 | 177 | ST-177 | ERP000129 | ERR024461 |
| 124 | starling45 | UK | 2007 | starling | 1,603,669 | 112 | 45 | ST-45 | ERP000129 | ERR024464 |
| 125 | starling1020 | UK | 2007 | starling | 1,578,916 | 61 | 1020 | ST-682 | ERP000129 | ERR024465 |
| 126 | goose1033 | UK | 2007 | goose | 1,663,834 | 149 | 1033 | ST-1034 | ERP000129 | ERR024466 |
| 128 | goose137 | UK | 2007 | goose | 1,600,143 | 53 | 137 | ST-45 | ERP000129 | ERR024468 |
| 129 | goose696 | UK | 2007 | goose | 1,561,449 | 143 | 696 | ST-1332 | ERP000129 | ERR024469 |
| 130 | duck702 | UK | 2007 | duck | 1,669,953 | 108 | 702 | ST-702 | ERP000129 | ERR024470 |
| 131 | duck45 | UK | 2007 | duck | 1,616,162 | 60 | 45 | ST-45 | ERP000129 | ERR024462 |
